# Is a long hygroscopic awn an advantage for *Themeda triandra* in drier areas?

**DOI:** 10.1101/2020.04.29.068049

**Authors:** Craig D Morris

## Abstract

*Themeda triandra* has bigeniculate hygroscopic lemma seed awns that twist when wet and drying, thereby transporting the caryopsis across the soil surface into suitable germination microsites. The prediction that awns would be longer in drier grassland and have greater motility to enable them to move quickly and further to find scarce germination sites was tested in KwaZulu-Natal. Awns (n = 100) were collected from 16 sites across a mean annual precipitation gradient (575-1223 mm), ranging from 271-2097 m a.s.l. The daily movement of hydrated long and short awns (n = 10) across blotting paper was tracked for five days, and the rotational speed of anchored awns was measured. Awn length varied considerably (mean: 41.4-63.2 mm; sd: 3.44-8.99) but tended to increase (r = 0.426, p = 0.099) not decline, with increasing MAP. Awn length was unrelated to elevation, temperature and aridity indices. Long awns rotated at the same rate (2 min 48 sec) but moved twice as fast (46.3 vs. 22.1 mm day^-1^) and much further (maximum: 82.1 vs. 38.6 mm day^-1^) than short awns. Whether moisture limits awn development, the benefit of longer awns to negotiate densely tufted mesic grassland, and the multifunctionality of awns require investigation.

---

Plants have developed diverse strategies to disperse their seeds, including evolving morphological structures that aid their movement (Howe and Smallwood 1982). In grasses, a common structure that enables autochory, or self-dispersal (van Rheede van Oudtshoorn and van Rooyen 1999), is a hygroscopic, twisted lemma seed awn that untwists when wet and coils again upon drying, thereby rotating the awn (Darwin 1877; Elbaum and Abraham 2104; Masrahi and Shaye 2017). A change in humidity of only 30% is required to induce movement in hygroscopic awns (Sindel et al. 1993). Hygroactivated rotating awns could drill the seed into the ground, and it has long been believed that the primary function of the hygroscopic awn is to assist in the burial of the seed (Darwin 1876; Pentz 1955; Zacharias 1990; O’Connor 1997; Snyman et al. 2013). However, Peart (1979) and Adams and Tainton (1990) found no evidence that awns frequently bury seeds but they did confirm that hygroscopic awns do at least move seeds across the soil, thereby increasing their chance of encountering and being orientated into suitable microsites for germination, such into cracks and crevices or under surface obstacles like litter and stones (Stamp 1989).

*Themeda triandra*, a palatable and productive tufted grass abundant in southern and East Africa as well as in Australia and parts of southern Asia (Snyman et al. 2013), has a single apical bigeniculate (twice-sharply bent) awn attached to the upper lemma of a callused seed (Cavanagh et al. 2019). Such a twisted geniculate lemma awn has evolved independently at least five times in various grass lineages, including in the subfamily Arundinoideae to which *T. triandra* belongs (Teisher et al. 2017). The active, basal twisted section of the awn extends to just past the last, most pronounced bend (‘knee’) whereas the tip section is passive, providing a lever for movement (Snyman et al. 2013; Masrahi and Shaye 2017). Awns vary in length from about 30 to 70 mm. Longer awns have been shown to bury seed of *Hyparrhenia diplandra* deeper than short awns in a West African savanna (Garnier and Dajoz 2001) whereas having a long active as well as passive section of an awn should confer greater torque and leverage for movement (Johnson and Baruch 2014).

This study aimed to test a hypothesis regarding the potential benefit of long awns that arose out of observations of genetically related variation in awn length of *T. triandra* along an aridity gradient in Australia. Diploid *T. triandra* plants with shorter awns occur along the wetter coastal regions whereas tetraploid plants with longer awns are found in the more arid interior (Godfree et al. 2017). They suggested that long awns are essential in drier areas to move seed as far and as rapidly as possible to find suitable microsites for germination after rain but that shorter awns should suffice in wetter areas where precipitation is higher and more predictable (Godfree et al. 2017). Because such variation in awn length has not been examined for *T. triandra* populations in South Africa, I tested elements of this hypothesis to assess whether (1) awns get longer with declining rainfall, and (2) long awns move faster and further than short awns on a given surface. The rotational speed and allometry of long and short awns were also examined to explain differences in motility.

Awns of *T. triandra* (n = 100) were collected in mid- to late summer from 16 sites in KwaZulu-Natal ranging in altitude from 271-2097 m a.s.l. across a mean annual precipitation gradient (MAP) from 575-1223 mm. The site locations are listed in Figure 1. The length of each awn with its caryopsis removed was measured stretched straight (Godfree et al. 2017) to the nearest millimetre. Linear regression was used to assess the relation between mean awn length per site and MAP, elevation, and various temperature (mean annual temperature and mean maximum temperature in February) and aridity indices (mean annual evaporation and plant available water) obtained from Schulze (2007).

**Figure 1:**
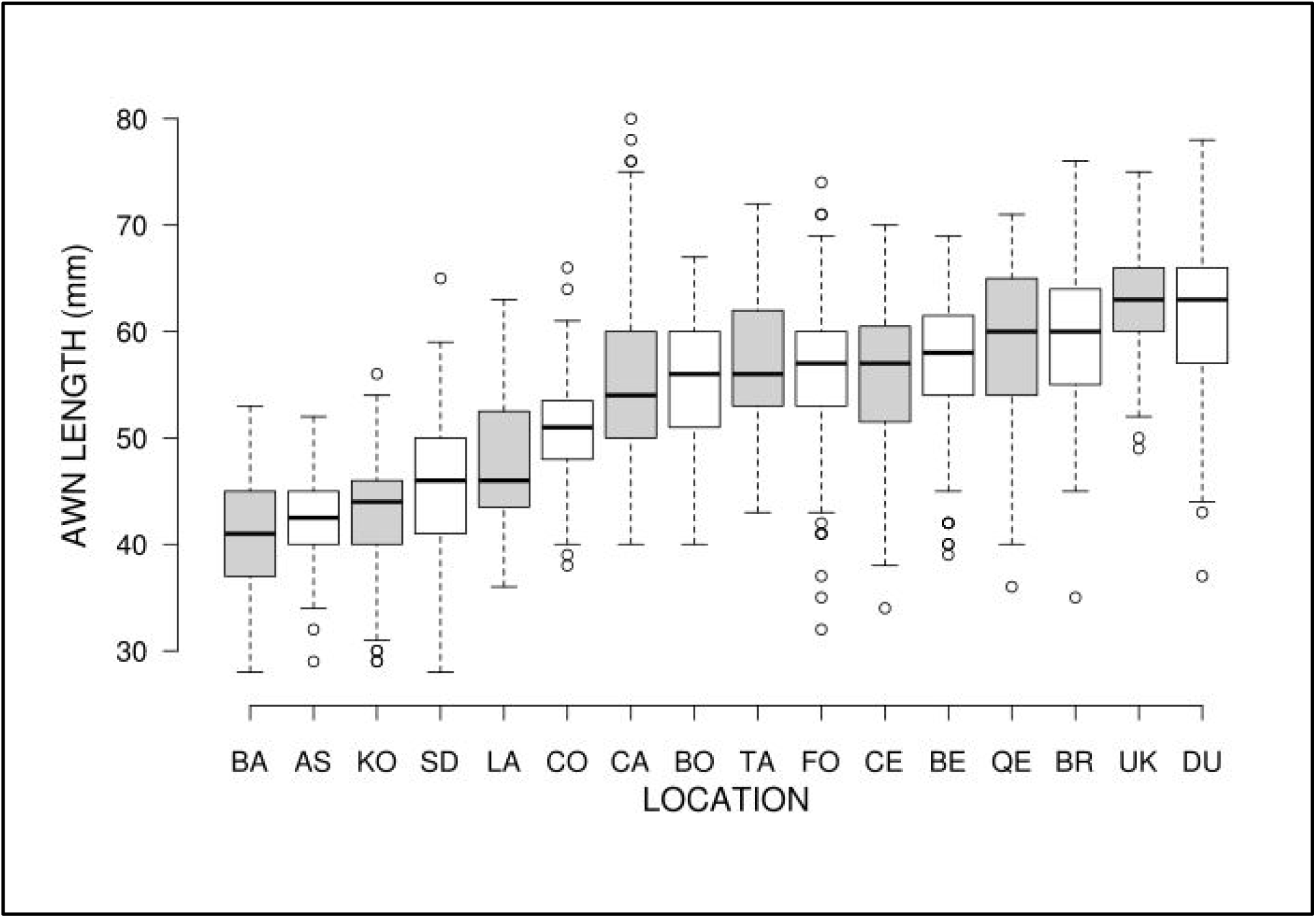
Median (horizontal bar) and interquartile range (boxes) of the length of hygroscopic awns of *Themeda triandra* collected from locations in KwaZulu-Natal (n = 100 per site). Locations are: B = Bartlow Research Farm; AS = Ashburton; KO = Kokstad Research Farm; LA = Ladysmith; CO = Colenso; CA = Cathedral Peak; BO = Boschberg farm; TA = Table Mountain; FO = Forth Nottingham; CE = Cedara Research Farm; BE = Beacon Hill; QE = Queen Elizabeth Park; BR = Brotherton Burning Trial; UK = Ukulinga Research Farm; DU = Dundee Research Farm.

The daily movement after hydroactivation of ten long (70.1 ± 5.79 mm) and ten short (41.6 ± 5.22 mm) awns across blotting paper (Adams and Tainton 1990) in two plastic trays (450 x 300 x 30 mm) was tracked and measured over five days. Each awn was randomly allocated to a track separated by plastic rulers (300 mm long) and placed with their seed on a central starting point on Day 1. Awns were sprayed until saturated each day and the distance they moved from the starting point and the previous day’s position was measured each day. Means for daily movement rate (mm day^-1^), total distance moved (mm), and maximum daily movement (mm) were compared using t-tests.

The spiral basal and non-twisted distal sections of 50 awns (44-73 mm long) were measured to correlate the relation between total awn length and proportional changes in the active and passive section of the awns. The time to complete a full rotation after wetting of 10 pairs of long (67.7 ± 5.93 mm) and short (43.0 ± 4.03 mm) vertically aligned awns, anchored at their base (Darwin 1876), was measured and compared with a paired t-test.

Awns ranged in length from 28 to 80 mm (mean: 53.1 ± 9.48 mm) and varied considerably among (mean: 41.4 - 63.2 mm) and within (standard deviation: 3.44 - 8.99; CV%: 7.67 – 16.20) sites (Figure 1). Variability (sd) tended to increase weakly with mean length (r = 0.487; p = 0.0554). The longest awns were 52.6% longer than the shortest awns, far greater variability than the 15% difference in awn length between tetraploid and diploid *T. triandra* plants recorded in Australia (Godfree et al. 2017).

Awn length tended to get longer with increasing MAP, but not significantly so (r = 0.426; p = 0.0998): short awns were uncommon in the wettest sites (Figure 2). Awn length was unrelated to elevation (p = 0.505) as well as to temperature variables and aridity indices (p > 0.30). This finding is contrary to the prediction that *T. triandra* plants with longer awns should predominate on drier sites (Godfree et al. 2017). The weak positive influence of annual rainfall suggests moisture availability cannot be discounted as a factor influencing awn growth and length, but other unmeasured environmental factors could have also contributed to the observed variability in awn length within and among sites. Also intriguing is the pattern of variation about the fitted line for MAP, where variation in awn length among sites was larger at the low than the high end of the rainfall gradient. Higher interannual moisture variability could be the driver of the observed variation in awn length among drier sites. Moisture and nutrient limitations during fruit development can restrict awn growth (Vaughton and Ramsey 1998) and drought can shorten awns (Godfree et al. 2017); greenhouse trials would be useful for testing this.

**Figure 2:**
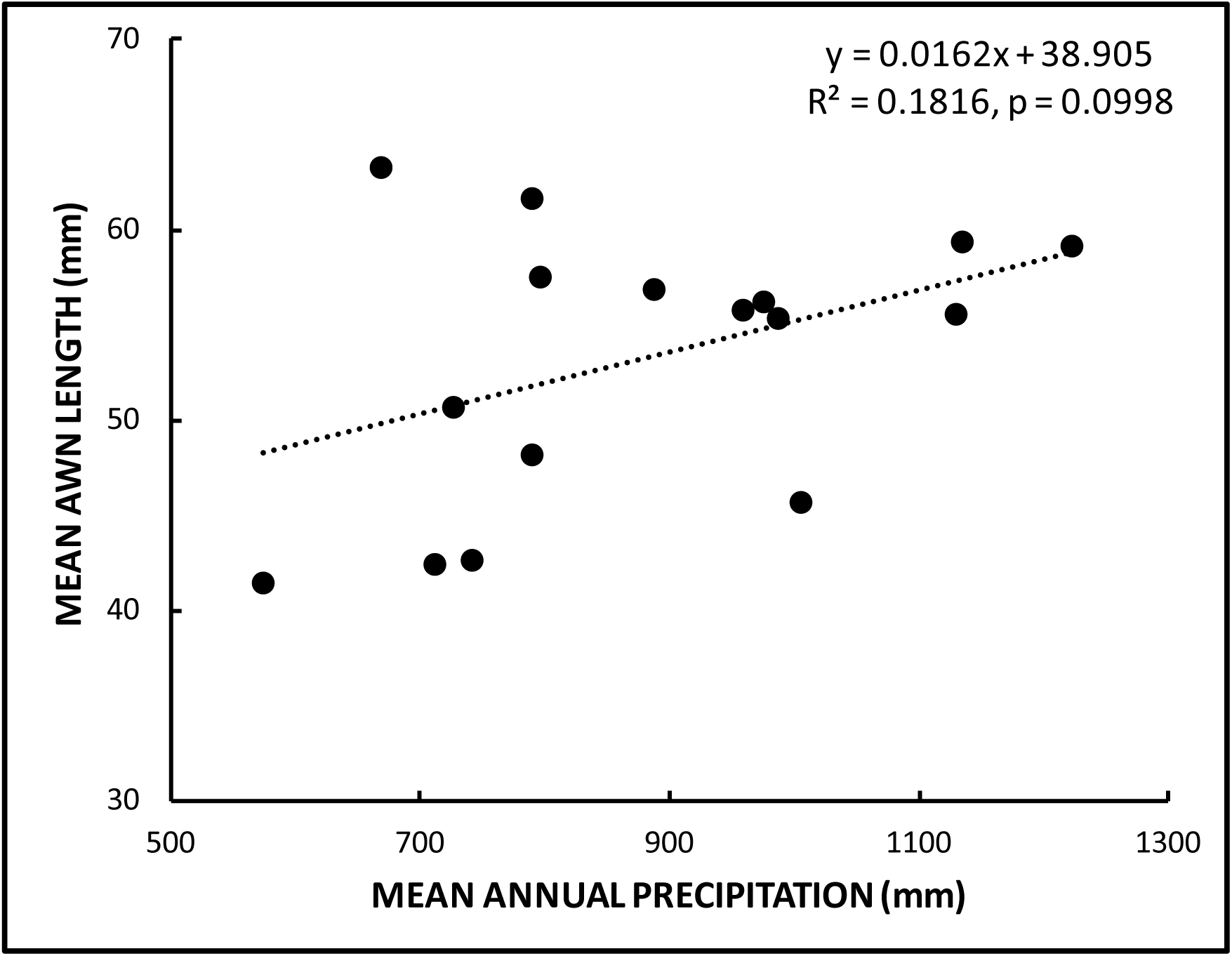
The linear relation (dotted line) between mean length of awns (n = 100 per site) of *Themeda triandra* and mean annual precipitation across 16 sites in KwaZulu-Natal, South Africa.

Genetics also probably play a role in determining potential awn length. The genome of *Themeda triandra* is a polyploidal complex that confers wide genetic variability between and even within populations (Liebendberg 1986; Fossey and Liebenberg 1987), making the species highly adaptable and morphologically plastic (Janse van Rensburg 2006). Garnier and Dajoz (2001) provide evidence that awn length is heritable for *Hyparrhenia diplandra* while Godfree et al. (2017) demonstrated that tetraploid *T. triandra* plants have consistently longer awns than diploid plants. The degree of polyploidy and the extent to which genome site influenced awn length were not determined for populations in my study. However, plants from all sites were likely to be tetraploids rather than diploids because the latter are more common on the coast in South Africa whereas plants in the interior and uplands of KwaZulu-Natal, where study sites were located, appear to be exclusively tetraploid (Liebenberg 1986), potentially with longer awns than coastal plants.

Irrespective of their determinants, longer awns were markedly more mobile than short awns (Figure 3). Long awns on average moved more than twice as fast (46.3 vs. 22.1 mm day^-1^) than short awns (t_18_ = 5.733, p < 0.0001), travelling a total of 121 mm further over five days (231.3 vs. 110.4 mm). The mean maximum daily distance moved from their source was also substantially higher (t_18_ = 3.177, p = 0.0026) for long than short awns (108.86 vs 56.81 mm) indicating their potential to disperse furthest from the parent plant. The width of the tray (300 mm) did restrict the maximum distance an awn could move from its origin. Awns did not travel consistently in one direction and some returned towards the start, resulting in a final net dispersal distance of 79.44 mm for long awns and 42.91 mm for short awns (t_18_ = 1.893, p = 0.0373) over five days. There are few data to compare these distances to. Drizin (2013) found that awned prairie grasses travelled less than 10 mm, unaffected by their length, whereas Stamp (1989) observed that the hygroscopic awn of the grassland forb, *Erodium moschatum* (Geraniaceae), can move seeds up to 70 mm across the soil surface, but not always away from the source plant. Wind can also assist hygroscopic awns of *Cymbopogon* spp. to disperse up 200 – 300 mm away from the mother plant (Ahmad et al. 2000). The daily distances moved by long awns in this study were similar to the approximately 45 mm and 60 mm travelled by *T. triandra* awns (of unspecified length, n = 2) when hydrated and drying, respectively (Adams and Tainton 1990).

**Figure 3:**
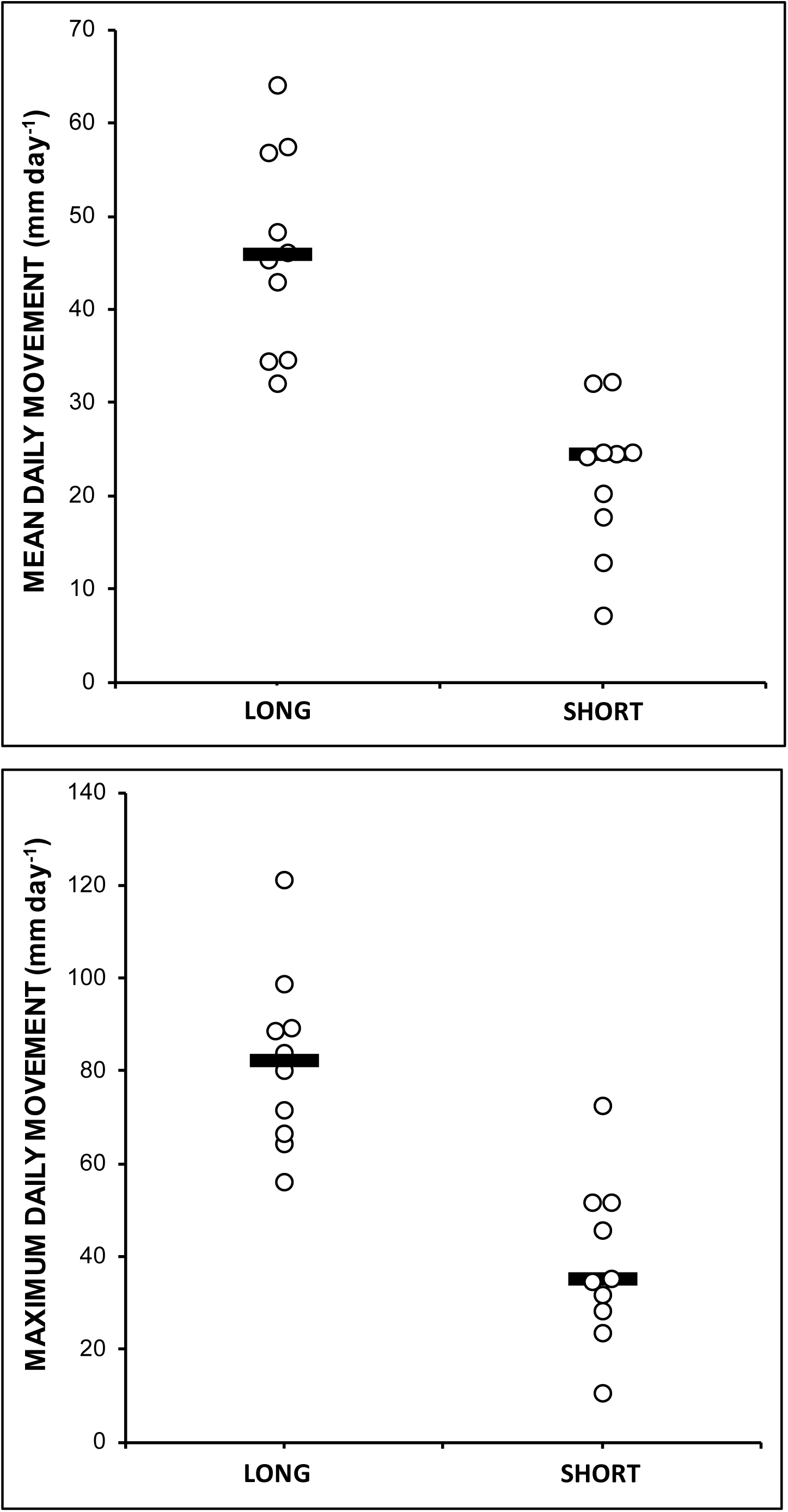
Mean (n = 10) daily movement and mean maximum distance moved per day for long and short awns of *Themeda triandra*. The horizontal bar is the median.

The morphological mechanism enabling long awns to move further than short awns was illuminated by examining their rotational movement and allometry. The basal helical section of the awn of *T. triandra* that provides the torque to power its movement was longest in longer awns (r = 0.926, p < 0.0001). The length of the passive lever (beyond the distal knee) also increased with awn length (r = 0.676,, p < 0.0001) but there was a disproportionate increase in the length of the active hygroscopic section of the awn (r = 0.504, p = 0.0002), which constituted just over 51% of the awn in short awns but more than 60% in long awns. However, when hydrated, long and short awns turned at the same rate (t_9_ = 0.107, p = 0.9168) taking on average 2 min 48 ± 15.2 sec (1 min 22 sec – 4 min 55 sec) to complete a full rotation. This time is in line with what Darwin (1876) found for the feathered awn of *Stipa pennata* (2 min 30 sec) and Silberbauer-Gottsberger (1984) measured for various grasses with hygroscopic awns (2-4 min) in the Cerrado of Brazil.

The reason why long awns did not twist faster than short awns is clear from Hooke’s law of angular momentum for helical torsion springs (Du and Xie 2013). The oblique helically arranged microfibrils in the inner water-absorbent cellular layer provides the torque in a hygroactivated coiled awn, which effectively acts a helical spring under tension (Elbaum and Abraham 2014; Masrahi and Shaye 2017). The length of the spring, in this case the hygroscopic portion of the awn, is not a determinant of the torque (force) exerted by the helical spring, which is only a mathematical function of the product of the inherent properties of the spring (the torsion coefficient) and how tightly it is coiled (the angle of twist). Longer active and passive sections do, however, increase the effective lever length and thus leverage exerted on the ground by an awn which pivots around a point (Johnson and Baruch 2014). In the awn of *T. triandra* the primary pivot is provided by the pronounced distal, sharply bended knee. The positive role of awn length in determining leverage thus explains why long awns move faster and further across the ground than short awns.

Hygroscopic awns, with their complex cellular morphology (see Elbaum and Abraham 2014) are probably metabolically expensive to construct, especially at the end of winter when resources are scare (Adams and Tainton 1990). Therefore, as Charles Darwin and his son believed, they likely evolved to serve an ecological purpose (Darwin 1876). Several studies (cited in Peart 1979 and Garnier and Dajoz 2001) have demonstrated that hygroscopic awns can actively burying seed. For example, long hygroscopic awns are most successful at burying seed deep enough to escape intense fires (Garnier and Dajoz 2001). Further studies on the potential role that the awn of *T. triandra* plays in seed burial should account for soil type because Peart and Clifford (1987) found grass species with hygroscopic and other active awns predominantly occur on structured, clay-rich soils with vertical cracks or loose, crumbly or broken surfaces that provide good seed burial sites. What is unequivocal, however, from my study is that awns do transport seed across a surface, with long awns being twice as effective than short awns. My study did refute the hypothesis that long awns would be favoured by aridity but did not provide any clear alternative explanation of what environmental factors influence the length of awns of *T. triandra* in KwaZulu-Natal. The tendency for the wettest sites to have long awns might reflect more favourable growing conditions there. An alternative or complementary explanation that requires testing is that longer awns might better promote the peregrinations of propagules though the densely-tufted landscape of a mesic grasslands where suitable sites to germinate in bare ground far from competitive mature tufts could be scarce (Everson 1994).

This study has conclusively established the role of awn length in motility for *T. triandra*. Hygroscopic awns could, however, also serve several other functions (Peart 1979) apart from transport and burial. Awns have been shown to render the seed heavy and cumbersome for ants to transport (Schöning et al. 2004) but Everson et al. (2009) observed very high predation of the seeds of *T. triandra*, which were carried up to 14 m by foraging ants (Camponotus and Myrmicaria spp.) in montane grassland. Ants also simply remove the nutritious endosperm through a small hole or slit in any unburied caryopsis remaining on the soil surface (Adams and Tainton 1990). Awns still attached to the inflorescence do twist when sprayed (personal observation) and thus could help free the seed for dispersal after rain. During its fall, the awn can also act as a flight to orientate the seed (Peart 1981) so it lands with the callus downwards onto the soil (Lock and Milburn 1970; Hendricks 2003). If trapped on leaves in the canopy during its fall (Darwin 1876) or in a tuft or other debris on the soil, hygroactivated movement could also help free the awn and seed. How long awns act to promote such multifunctionally still requires much research.

## Acknowledgements

I am grateful to Debbie Jewitt and Janet Taylor who helped organise the collection of awns and to Colin Everson and Kevin Kirkman for their awn samples. Debbie Jewitt also provided the environmental data and Anita Morris assisted with the movement experiments.

